# Flow stimuli reveal ecologically-appropriate responses in mouse visual cortex

**DOI:** 10.1101/362459

**Authors:** Luciano Dyballa, Mahmood S. Hoseini, Maria C. Dadarlat, Steven W. Zucker, Michael P. Stryker

## Abstract

Assessments of the mouse visual system based on spatial frequency analysis imply that its visual capacity is low, with few neurons responding to spatial frequencies greater than 0.5 cycles/degree. However, visually-mediated behaviors, such as prey capture, suggest that the mouse visual system is more precise. We introduce a new stimulus class—visual flow patterns—that is more like what the mouse would encounter in the natural world than are sine-wave gratings but is more tractable for analysis than are natural images. We used 128-site silicon microelectrodes to measure the simultaneous responses of single neurons in the primary visual cortex (V1) of alert mice. While holding temporal-frequency content fixed, we explored a class of drifting patterns of black or white dots that have energy only at higher spatial frequencies. These flow stimuli evoke strong visually-mediated responses well beyond those predicted by spatial frequency analysis. Flow responses predominate in higher spatial-frequency ranges (0.15–1.6 cycles/degree); many are orientation- or direction-selective; and flow responses of many neurons depend strongly on sign of contrast. Many cells exhibit distributed responses across our stimulus ensemble. Together, these results challenge conventional linear approaches to visual processing and expand our understanding of the mouse’s visual capacity to behaviorally-relevant ranges.

**Significance Statement:** The visual system of the mouse is now widely studied as a model for development and disease in humans. Studies of its primary visual cortex (V1) using conventional grating stimuli to construct linear-nonlinear receptive fields suggest that the mouse must have very poor vision. Using novel stimuli resembling the flow of images across the retina as the mouse moves through the grass, we find that most V1 neurons respond reliably to very much finer details of the visual scene than previously believed. Our findings suggest that the conventional notion of a unique receptive field does not capture the operation of the neural network in mouse V1.

## Introduction

The mouse has become a major model for studying vision because of the genetic, imaging, and molecular tools available [1]. Studies have revealed relationships between macroscopic states of the brain and activity in visual cortex (running *vs.* stationary [2,3], pupil size and activity [4,5], and visual interest (e.g., [5–7]). However, a basic conundrum has arisen: behaviorally, mice are capable of sharp, visually-mediated behaviors [8–10], such as accurate prey capture [11], but when assessed using standard assays, such as spatial frequency gratings (Fig. 1), the mouse appears to have very poor vision. Although orientation-selectivity has been found [12], receptive fields are large (typically ≥25 degrees^2^) when estimated by spike triggered averaging, and spatial frequency tuning is concentrated below 0.08 cycles/degree (cpd). While this motivates the use of gratings at 0.04 cpd in experiments, it raises the question: How does the visual system perform so exquisitely in natural tasks?

We show here that ecologically-relevant stimuli can exercise mouse visual cortex in novel and manifold ways. While plaids [13,14] and random-dot kine-matograms [15,16] are a step beyond gratings, the leap to natural images (e.g., [17]) is more common (e.g., [18,19]). However, natural images are difficult to obtain [20], difficult to control parametrically, and difficult to analyze beyond second-order [21].

**Figure 1:**
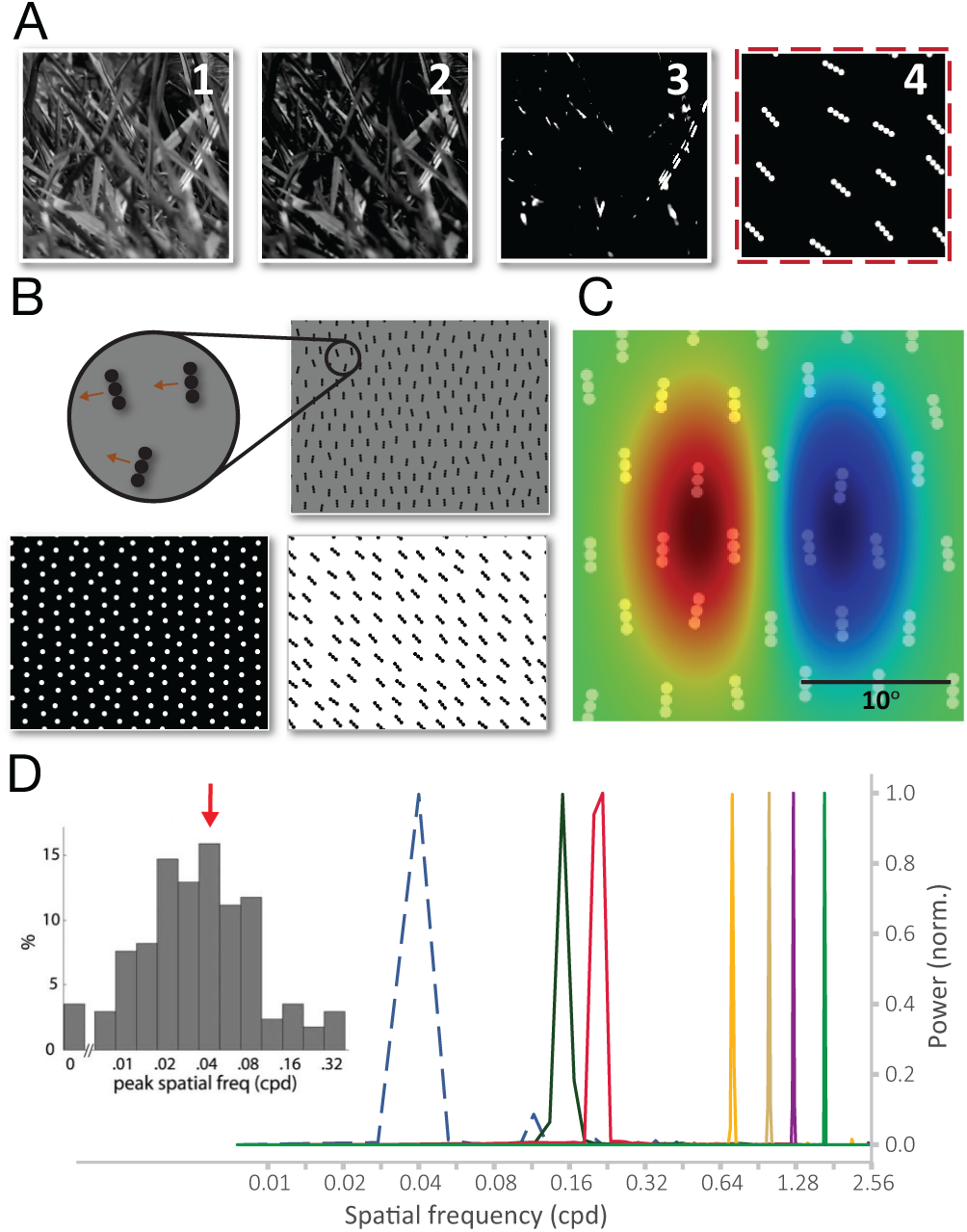
Introducing flow stimuli. ***A***, Ecological motivation: Working with a single frame (panels 1–3), a grassy patch is modified to emphasize salient higher contrasts. Our abstraction (the flow field in panel 4) approximates this with a binary pattern of random, oriented dotted segments. ***B***, We generalize to flow fields consisting of dotted segments of different lengths, emphasizing two geometries (oriented [3 or 4 aligned dots] or non-oriented [single dots]), two contrast polarities (positive or negative), different contrast magnitudes, and various sizes. The full flow stimulus is a movie of one such flow field, drifting across the screen with small random perturbations to suppress rigidity (see *Materials and methods*). ***C***, Flows are inconsistent with classical filtering views of V1. A Gabor receptive field at 0.04 cpd superimposed onto the 3-dot flow whose energy peaks at 0.24 cpd (top right example in ***B***), for comparison. ***D***, The 1-D discrete Fourier transforms (single-sided) of the flows utilized in our experiments (peaks at 0.15, 0.24, 0.7, 1.0, 1.25, and cpd) have power well beyond 0.04 cpd (dashed curve), which is the spatial frequency previously reported as optimal for cells in mouse V1 (cf. inset, from [12]). To compare stimuli, each spectrum is normalized by the power at the peak frequency (2D spectra in *SI* Fig. S14-S15).

For a mouse running through a field, the visual display is like a ‘waterfall’ of illuminated material flowing past, with bright or dark oriented segments arising from complex photometric events (Fig. 1A)” [22]. This visual metaphor motivates our stimuli. We approximate such patterns with a class of visual flows comprised of dots. These are more natural than drifting gratings but can be parametrically controlled in their orientation (content and angle), spatial frequency, and direction of motion. We call them flows because, intuitively, they consist of a field of particles (either dots or dotted line-segments) dropped into a ‘flowing river’. More formally, each dot is displaced along a vector field in space and time and follows a dynamical system [23]. When the orientation structure is removed, the flows reduce to random-dot kinematograms; when the temporal structure is removed, the flows reduce to static Glass patterns [24]. Thus they are rich in geometry and, for humans, the perception of such flows differs from strictly aligned patterns [25,26]. Parametric variations in orientation, direction, etc., define an ensemble of stimuli.

We here explore activity in mouse V1 in response to the flow ensemble. In many cases flow stimuli elicit more vigorous responses than drifting gratings, particularly at high spatial frequencies 3–5 octaves above 0.04 cpd. Some V1 neurons are classical, resembling feature detectors, while others exhibit a mixed selectivity rarely reported in early visual cortex. The rich ensemble of selectivities in V1 may equip the mouse to behave in the natural world.

## Results

### Analysis of stimulus selectivity in V1

#### Cells in V1 have diverse preferred stimuli

We developed an ensemble of stimuli including drifting gratings, single dot flows (random dot kinematograms), and oriented flows where each element consists of 3 or 4 dots (see Methods). The stimuli had either positive contrast (bright dots) or negative contrast (dark dots). Activity is plotted as an array of peristimulus time histograms (PSTHs) and tuning curves for each unit, to facilitate a quick assessment of the different “dimensions” of a cell’s response. Experiments were conducted in two cohorts, the first with grating stimuli at 0.04 cpd and both grating and flow stimuli at 0.15 and 0.24 cpd, and the second cohort with grating stimuli at 0.04 cpd and both grating and flow stimuli at 0.7, 1.0, 1.25, and 1.6 cpd. All stimuli in both cohorts had a fixed temporal frequency of 4 Hz.

We begin with example cells from cohort 1. The first one (Fig. 2A) has the response profile one would expect for a *simple cell* in V1. It responds almost exclusively to low-frequency gratings; the PSTHs for high-frequency gratings and for flows (both one dot and three-dot elements) remain virtually at baseline. Its spike triggered average (STA) depicts a classical receptive field, consistent with the frequency response, and it is well-tuned for orientation. But such cells were relatively rare in our experiments (discussed below). Another example (Fig. 2B) exhibits a weak response to gratings and a stronger response to flows. The STA, which would predict a strong response to low-frequency gratings, completely fails to predict this response profile. Finally, many cells are multidimensional (Fig. 2C): they respond well to several stimuli from the ensemble, including gratings and flows at multiple spatial frequencies. Note the diversity in the temporal response profile: a periodic (often interpreted as linear) response to gratings at low spatial frequency; a sustained (interpreted as nonlinear) response to gratings at higher frequencies; and a transient burst of activity to positive, oriented flows. It would be inappropriate to label this cell a classi cal feature detector. The STA again does not predict the response profile, and the PSTHs reveal different tuning widths, different first-spike latencies, as well as linear *vs.* non-linear and transient *vs.* sustained responses.

**Figure 2:**
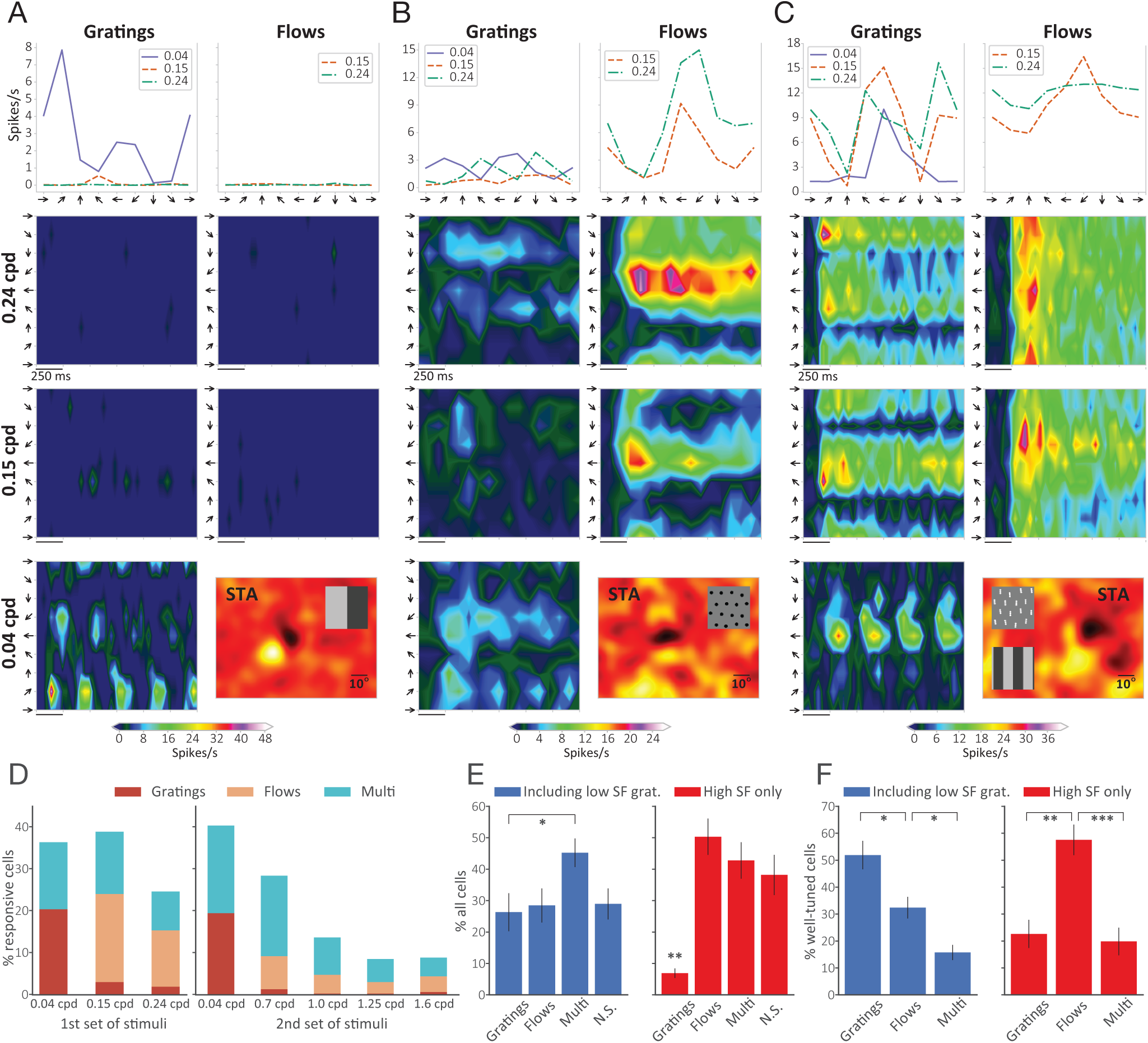
Variety of responses in V1. ***A–C***, Tuning curves and PSTHs of three example cells in response to drifting gratings and flows at 0.04, 0.15, and 0.24 cpd in 8 equally-spaced directions of motion. Time axis in histograms encompasses an entire period of stimulus presentation (1.5 s). Insets in STAs show, at the same scale, stimuli that produced the most significant responses. ***A***, Cell responding to low-frequency gratings only. Bin size 34 ms. ***B***, Cell responding preferentially to single-dot flows with negative contrast. Bin size 83 ms. ***C***, Cell responding strongly to both oriented (3 dots), positive flows and gratings (at both high and low spatial frequencies). Bin size 46 ms. ***D***, Distribution of optimal spatial frequency in terms of proportion of cells significantly responding to at least one of the stimuli. In the group of experiments using the first set of stimuli (left panel, 0.04–0.24 cpd, n = 357 cells, 3 animals), the majority of cells fired more strongly for stimuli at 0.15 cpd, followed closely by 0.04 cpd. For the second set of stimuli (right panel, 0.04–1.6 cpd, n = 256 cells, 3 animals) there was an overwhelming preference for 0.04 cpd, although more than half the cells had optimal spatial frequency in the range 0.7–1.6 cpd. ***E***, Distribution of preferred stimulus among all cells. When low-frequency gratings (0.04 cpd) are included among the stimuli (left panel), the majority of cells respond equally well to both classes (“Multi”), followed by only flows and only gratings; 29% of the cells were not significantly responsive (“N.S.”) to any of the stimuli displayed (n = 1026 cells; 10 experiments, 6 animals). When we do not include low-frequency gratings, thereby limiting the comparison to flows and gratings with similar spatial frequencies, there is a significant preference for flows only and for both over gratings only. Comparison of the left and right panels reveals that approximately 20% of cells preferred low-frequency gratings. When we recompute stimulus preference considering only stimuli with comparable spatial frequencies, most cells that preferred low-frequency gratings now either prefer none of the high-frequency stimuli, or significantly prefer flows over high-frequency gratings, given that the fraction that prefers both remains essentially constant in the two scenarios. Error bars represent s.e.m. ***F***, Distribution of preferred stimulus among well-tuned cells (i.e., those with OSI > 0.5 or DSI > 0.5), n = 295 cells (left), 241 cells (right); 8 experiments, 4 animals. Here, notice that most of the cells responding to orientation and/or direction will fire more strongly to low-frequency gratings; the right panel reveals, however, that the fraction of cells well-tuned to flows is just as large. And, similarly to ***E***, many of the well-tuned cells preferring 0.04 cpd gratings prefer flows to gratings of comparable spatial frequency. Error bars represent s.e.m.

#### Responses to optimal flows span a wide range of spatial frequencies

To quantify this diversity at the population level, we relaxed the notion of a unique preferred stimulus for a cell to allow for multiple possible preferences, according to the following definitions. While this leads to a crude classification of cell types, we stress that it is merely a set of labels for discussion; the underlying complexity remains in the PSTHs.

An individual stimulus is *significant* for a particular cell if the average firing rate for that stimulus is significantly higher than that for its preceding interstimulus interval (Mann-Whitney test). A cell prefers a *stimulus class* (e.g., flows or gratings) if at least one variation of that class (spatial frequency, geometry, or contrast polarity) is significant and has average peak firing rate significantly higher than the peak firing rates of all significant variations of the other class (Kruskal-Wallis rank-sum test, Conover-Iman post-hoc, Bonferroni correction, p < 0.05). When there is no preferred stimulus class but there are significant stimuli in both classes, we classify the cell as multi-class, or simply multi. Thus the preferred stimulus class, or *type* of a cell, is one of grating, flow, multi, non-selective. By this classification, the cell in Fig. 2A would be classified as a grating cell; Fig. 2B would be a flow cell; and Fig. 2C would be a multi cell.

Once each cell’s type, or preferred stimulus class, has been determined, its *preferred spatial frequency* can be defined as the one with highest average firing rate among all significant variations of the preferred class (or classes, when cells are labeled MULTI).

We plot the proportion of preferred types at each preferred frequency in Fig. 2D; the two separate plots denote units from experimental cohort 1 (0.04–0.24 cpd, n = 357 cells, 3 animals) and cohort 2 (0.04–1.6 cpd, n = 256 cells, 3 animals), respectively. Note the predominance of gratings among cells at the lowest frequency, replicating the inset in Fig. 1D, and the predominance of flow and multi types at the higher frequencies.

We now examine the distribution of preferred *types* in two different ways, either including or not including the responses to low-frequency gratings. This is necessary, since the performance measure is a simple spike statistic that is easily dominated by the gratings. First, when low-frequency gratings are included among the stimuli, by the above definitions 45% of the cells respond equally well (i.e. are of the multi type); 28% are flow cells; 26% prefer gratings; and 29% of the cells were not significantly responsive to any of the stimuli displayed (see Fig. 2E, blue). When low-frequency gratings are not included, so that the comparison is among flows and gratings at the same spatial frequencies, responses favoring flow (50%) and multi (43%) predominate over those to gratings (7%) (see Fig. 2E, red). The difference between these two plots comes from a more detailed analysis: the cells responding strongly to 0.04 cpd can be divided into roughly two subgroups: one that has no significant response other than to low-frequency gratings and another that also responds well to flows (or, in fewer cases, to both flows and high-frequency gratings). These plots include all cells. A similar distinction obtains when only cells well-tuned to orientation (OSI *>* 0.5) or direction (DS *>* 0.5) are considered (Fig. 2F).

To summarize, among cells with significant preference for flows or both flows and gratings, responses were distributed across all spatial frequencies explored. For “classical” cells (those that significantly preferred gratings to flows) there is a clear preference for 0.04 cpd with a distribution in accordance with [12] (see Fig. 1D). Curiously, some cells that are well-tuned to low-frequency gratings are also well-tuned to flows with higher spatial frequency, albeit usually with lower firing rates. Nevertheless, many of these cells have higher firing rates to flows than to gratings of similar spatial frequency, showing that there is some aspect of the flow stimulus that strongly excites these cells despite the fact that the flow elements would not excite the filter predicted by these cells’ STAs. Supplemental Materials Fig. S1–S13 show plots of responses to the entire stimulus ensemble for these and other cells.

#### Cells remain well-tuned at high spatial frequencies

Since higher firing rates do not necessarily imply high orientation- or direction-selectivity, and since a cell might retain its selectivity at several spatial frequencies (SFs), we investigated the fraction of well-tuned cells (OSI > 0.5 or DSI > 0.5) across spatial frequencies regardless of preferred stimulus (Fig. 3B). This is an estimate of the probability of a cell significantly responsive to a certain SF being well-tuned.

**Figure 3:**
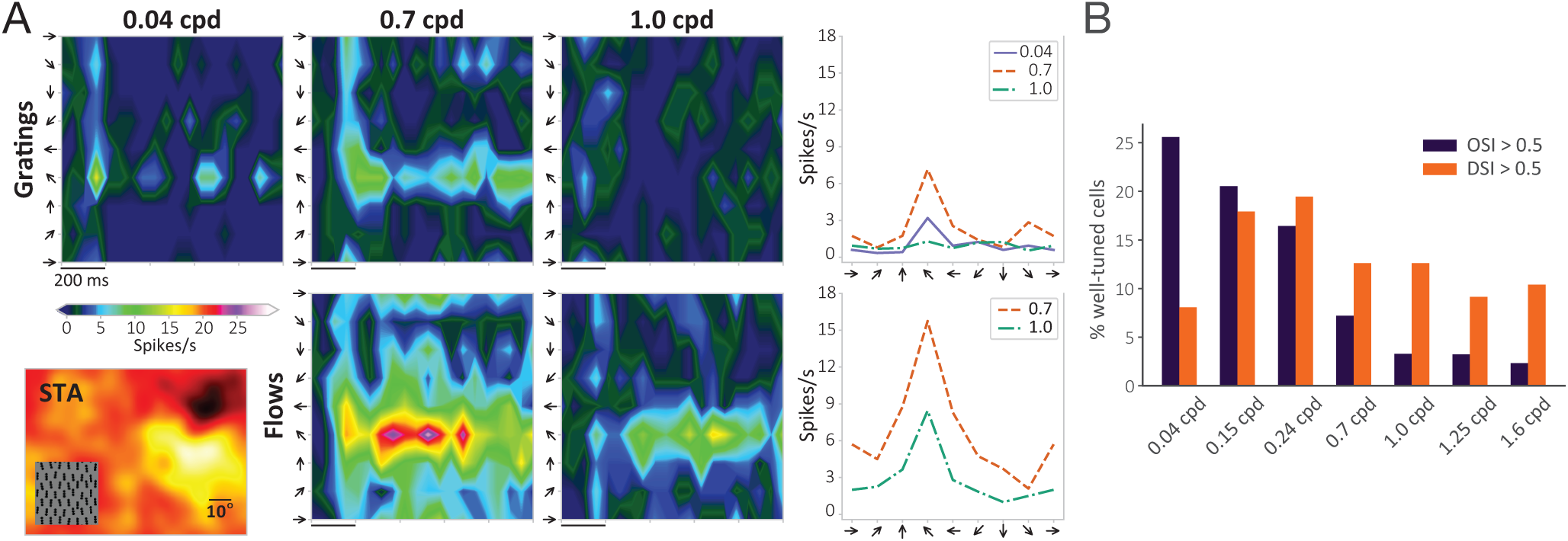
Cells remain highly selective at higher spatial frequencies. ***A***, Example of cell exhibiting a stronger response to oriented, negative flows at 0.7 and 1.0 cpd when compared to gratings at various spatial frequencies. Bin size 47 ms. ***B***, Overall proportion of well-tuned cells among cells significantly responsive to each spatial frequency (Mann-Whitney test, p < 0.05), irrespective of stimulus class. Sample sizes: 0.04 cpd (n = 508), 10 experiments, 6 animals; 0.15 cpd (n = 385), 0.24 cpd (n = 365): 5 experiments, 3 animals; 0.7 cpd (n = 214), 1.0 cpd (n = 214), 1.25 cpd (n = 186), 1.6 cpd (n = 173): 5 experiments, 3 animals.

There are many cells well-tuned to direction and/or orientation at all SFs. Cells with high orientation selectivity tend to prefer stimuli in the 0.04–0.24 cpd range. The direction-selective cells seem to be more uniformly distributed across SFs, with a preference for intermediate SFs (0.15–0.24 cpd).

#### Higher stimulus selectivity in superficial and deep layers

To further characterize how the response profile of multi cells is distributed across stimulus variations, we extend the concept of selectivity indices such as OSI and DSI (e.g., [12]) to compare pairs of stimulus classes. A stimulus selectivity index (SSI) is thus defined for a pair of classes (e.g., flows *vs.* gratings, or 1-dot flows *vs.* 3-dot flows) as (*R*_max_ − *R*_min_)/(*R*_max_), where *R*_max_ (*R*_min_) is the average peak firing rate (FR) of the stimulus with the higher (lower) FR in the pair. Essentially, it measures the difference in FR between two stimuli, relative to the one with highest FR. E.g., an SSI of 0.2 means the FR for the less preferred stimulus is 20% lower than that for the preferred one. When comparing stimulus classes for which there are possibly several stimulus variations in each class, we take the variation that elicited the highest response in each one. Note that the SSI for a cell population assesses how well those cells’ responses can be used to differentiate between two stimuli, regardless of which one is the preferred one.

Cells responsive to both flows and high-frequency gratings were found in all cortical layers. Cells in layer 2/3 had significantly higher values of SSI than all other layers for differentiating flows from gratings (p < 10^-3^, p = 10^-6^, and p = 10^-4^ for layers 4, 5, and 6, respectively), while cells in layer 5 had significantly lower SSI than layers 2/3 (p < 10^-4^) and 6 (p < 0.05) when differentiating between flows with opposite contrast polarities, and lower than layer 2/3 (p < 0.005) when differentiating oriented from non-oriented flows (Fig. 4). The same trends were found when only broad-spiking cells (putative excitatory, see [12]) were considered. Thus, speculatively, cells in the superficial layers could have higher selectivity, while cells in layer 5 could be more invariant to geometry, length, and contrast. This may be related to [12], in which it was reported that layer 5 cells were significantly less linear than cells in other layers.

**Figure 4:**
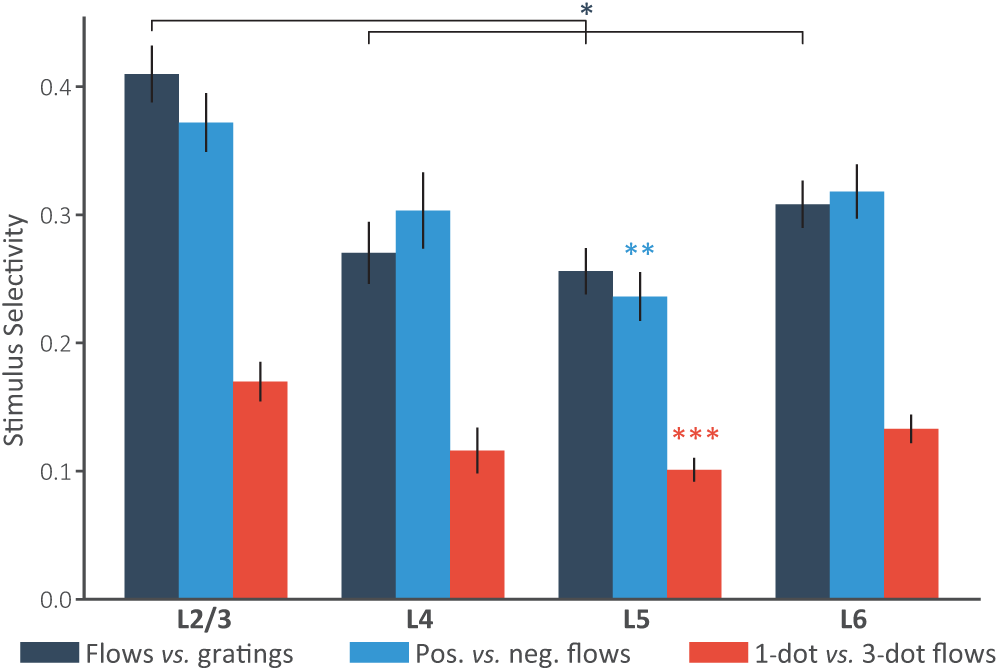
Cells in different layers have distinct selectivity toward different stimulus classes, as measured by a stimulus selectivity index (SSI, see text). Error bars represent s.e.m.

#### Preference among different variations of flow stimuli goes beyond differences in spatial frequency

Among cells that responded significantly to flows, we also compared the average proportion of cells that significantly preferred oriented (3 dots) *vs.* non-oriented flow patterns (single dots) (Fig. 5A). Analysis of the entire population across different experiments does not reveal any particular preference, with the vast majority responding to both geometries. However, if analysis is restricted to those cells well-tuned to direction and/or orientation, the preference for a specific flow geometry—be it oriented or non-oriented—increases markedly. In particular, there is an overall preference for the oriented patterns.

**Figure 5:**
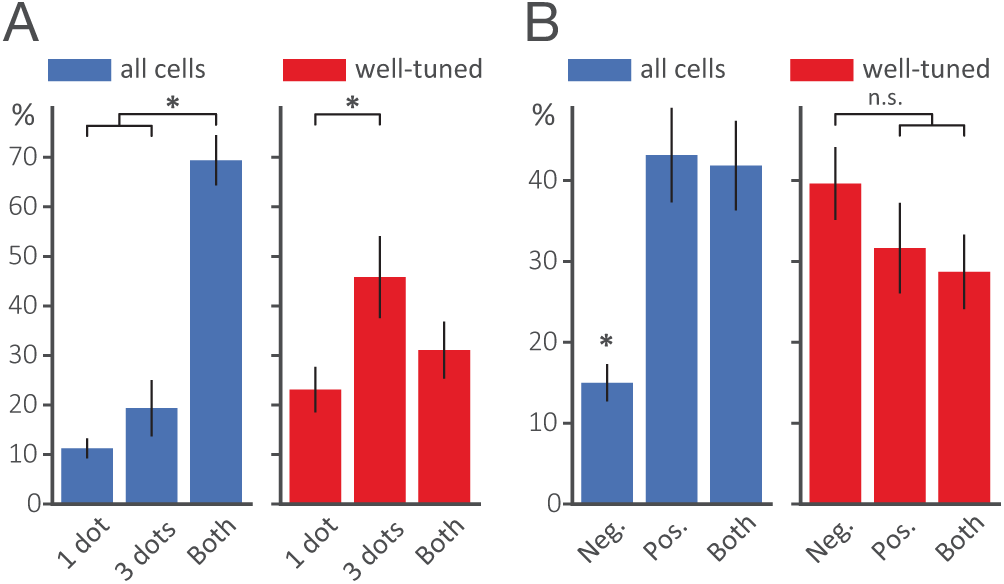
Preference over flow stimuli variations. Percentages refer to the population of cells that had significant response to at least one flow variation. All cells: n = 667, 10 experiments, 6 animals; well-tuned cells: n = 187, 8 experiments, 4 animals (applies to panels ***A*** and ***B***). Error bars represent s.e.m. (*, p < 0.001). ***A***, Flow geometry preference. Among all cells responding significantly to flows, most showed no significant prefernce for either type. Among well-tuned cells, oriented flows were preferred over non-oriented flows. ***B***, Flow contrast polarity preference. Among all cells significantly responding to flows, positive polarity was preferred. The population of well-tuned cells showed no overall preference for contrast polarity.

Fig. 5B shows that only a minority of the cells responding to flows prefer negative contrast (15%, on average). The vast majority prefer either positive contrast or respond significantly to both contrast polarities. This difference in preference disappears among cells that are well-tuned to direction and/or orientation.

## Discussion

### Receptive fields redux?

Cells are routinely classified as “simple” if they exhibit a linear response to moving bars or gratings [27] and as “complex” if the response is non-linear (over phase). However, these are operational definitions and depend on the stimuli. Arguments against such classical receptive field concepts are developing [28], and our results contribute to this. Unlike the traditional, optimal feature viewpoint, selectivity does not appear to be one-dimensional; cells can respond linearly to one part of the stimulus ensemble while being non-linear to others. In particular, we found many cells that exhibit a ‘linear’ response to low-frequency gratings while also responding vigorously at high spatial frequencies to some type of flow (Fig. S16 in *SI*).

The fact that linear methods (e.g., [29]) cannot explain such complex and varied responses to an ensemble of stimuli has several implications (Fig. S17-S18 in *SI*). First, it brings some of the feature variability seen in higher visual areas (e.g., [30,31] and references therein) down to V1. Second, since the receptive field is often taken as the signature of functionality, attempts to relate function to structure based on it (e.g., [32]) may attribute to ‘noise’ in connectivity genuine features important for neural responses. It follows that stimulus ensembles richer than low-frequency gratings are required to properly assess visual system function. Finally, it suggests that network computations, not individual features, should be the focus of investigation [33].

### Flow responses suggest network computations

At a small scale, responses to high-frequency flow stimuli (i.e., within a receptive field) are reminiscent of the sub-zones observed in the fly [34] and primate [35], each of which can be direction- or orientation-selective. And, as in the fly [36], many cells with a preference for flow stimuli are also contrast selective. At a large scale, our stimuli were displayed wide-field, so extra-classical effects may also be playing a role. While investigations of such contextual interactions in the mouse are just beginning, when stimuli were restricted to gratings [37] and bars [38], only suppressive effects were observed [39]. By contrast, in the primate the situation is much richer [40–42], and the arrangements of dots and bars in these studies are reminiscent of our flow stimuli. Despite the fact that the mouse lacks orientation columns [43], flow responses are remarkably consistent among the different species. Flow stimuli are also informative about the geometry of surrounding objects [44], though how these geometric, network computations are realized remains an open question.

## Materials and Methods

### Animal procedures

Experiments were performed on adult C57/BL6 mice (age 2–6 months) of either sex. The animals were maintained in the animal facility at the University of California–San Francisco and used in accordance with protocols approved by the University of California–San Francisco Institutional Animal Care and Use Committee. Preparation, extracellular recording, and single neuron analysis were generally performed as in [45]. See Supplementary Information for additional material and methods.

### Design of flow stimuli

The flow stimuli consist of local flow elements that move according to an underlying displacement field. The displacement field is defined as a vector in ℝ ^2^ (i.e., a magnitude and a direction) at each screen position. Each flow element consists of a linear arrangement of *n* adjacent dots, with *n* = 1 corresponding to a random-dot kinematogram, and *n* = 2, 3, 4 corresponding to oriented elements (cf. Fig. 1B). The stimulus density defines an integer screen lattice. Each flow element is dropped onto this lattice, and then its position is perturbed by a normally-distributed random variable. This destroys the impression of a perfectly regular grid of elements.

The displacement field is built on top of this, and is organized around a screen partition consisting of a grid of rectangular or hexagonal tiles, with a single displacement vector within each tile. The tiles provide controllable flexibility in the motion. A planar translation results from choosing all displacement vectors the same. To avoid the impression of such simple translations, each displacement vector is ‘jittered’ by sampling from a normal distribution, so that there is now both position jitter and motion jitter. More generally, one can develop more complex motions, either in geometry or in time, by varying the size of the tiles (discreteness) and by varying the displacement directions. For the experiments reported in this paper, a common mean and variance for all vectors in the field was used (see SI Methods for specific parameter values).

The movement of each flow element is made even more “lifelike” by controlling its acceleration with a steering force computed as the difference between the element’s desired and current directions of motion. This behavior is based on the boids from [46] and on the steering behavior described in [47]. Using this, the desired direction can be a function of both the underlying flow field and the proximity between elements; this guarantees that elements do not overlap one another, creating different geometries, densities, or sizes. The final force applied to each element is the resultant between the steering force and a repulsion force exerted by every other element within an allowed distance. The advantage of this approach is that the flow elements can make successive changes in direction as they drift through the flow field by following a smooth and continuous trajectory, without abrupt changes in direction or occlusion. They also wrap around the screen boundaries to preserve a constant number of patterns being shown at all times.

It remains to control the overall luminance and its changes, both for different stages in the trial and for the local dots. For some experiments, the screen luminance during the interstimulus period was set equal to the global average luminance of the stimulus used in the upcoming trial. This controls for responses that could be caused by the global change in the screen luminance only, and not by the actual moving stimuli. However, for this strategy there will still be a change in the luminance of the background when the trial starts. To control for this, i.e., for possible responses due to changes in background luminance, we also ran experiments with a constant gray background both during the flow trials and the interstimuli intervals. For dot-luminance effects, the diameter of each dot in a multi-dotted flow element was chosen such that the total area occupied by the element was the same as that of the single-dotted version of the stimulus with same spatial frequency. In any case, repeated experiments showed that there was little (if any) effect of these different luminance change variations. Code for generating the flow stimuli is available at https://github.com/zuckerlab/FlowStims.

## Acknowledgements

Supported by the Simons Collaboration on the Global Brain and the National Eye Institute grant R01-EY02874. M.P. Stryker is the recipient of the RPB Stein Innovation Award. We are grateful to Dr. Sotiris Masmanides of UCLA for supplying the microelectrodes through the NSF NeuroNex program.

